# Rad51 is a druggable target that sustains replication fork progression upon DNA replication stress

**DOI:** 10.1101/2022.03.25.485778

**Authors:** Sònia Feu, Fernando Unzueta, Amaia Ercilla, Montserrat Jaumot, Neus Agell

## Abstract

Solving the problems that replication forks encounter when synthesizing DNA is essential to prevent genomic instability. Besides their role in DNA repair in the G2 phase, several homologous recombination proteins, specifically Rad51, have prominent roles in the S phase. Using different cellular models, Rad51 has been shown not only to be present at ongoing and arrested replication forks but also to be involved in nascent DNA protection and replication fork restart. Through pharmacological inhibition, here we study the specific role of Rad51 in the S phase. Rad51 inhibition in non-transformed cell lines did not have a major effect on replication fork progression under non-perturbed conditions, but when the same cells were subjected to replication stress, Rad51 became necessary to maintain replication fork progression. Notably, the inhibition or depletion of Rad51 did not compromise fork integrity when subjected to hydroxyurea treatment. Rad51 inhibition also did not decrease the ability to restart, but rather compromised, fork progression during reinitiation. In agreement with the presence of basal replication stress in human colorectal cancer cells, Rad51 inhibition reduced replication fork speed in these cells and increased γH2Ax foci under control conditions. These alterations could have resulted from the reduced association of DNA polymerase α to chromatin, as observed when inhibiting Rad51. It may be possible to exploit the differential dependence of non-transformed cells versus colorectal cancer cells on Rad51 activity under basal conditions to design new therapies that specifically target cancer cells.

## Introduction

DNA replication is a moment of major genome vulnerability(1). In eukaryotes, DNA replication is initiated from multiple origins distributed along the chromosomes(2). Once fired, each origin generates two replication forks that move away in opposite directions led by separate replisomes. The replisomes are large protein complexes that take part in DNA replication and contain, as essential components, the Cdc45-MCM-GINS (CMG) helicase involved in unwinding parental DNA; the DNA polymerases α, δ, ε responsible for incorporating nucleotides; and DNA polymerase processivity factors(3). As the replisome progresses, it may encounter several problems that cause a functional uncoupling between polymerases and helicase, stalling replication forks and challenging cells with replication stress. These problems can be solved through error-free or error-prone mechanisms, the latter being one of the causes of genomic instability(4–6). Extrinsic factors, such as DNA alkylating or crosslinking agents, may cause problems with the replisome, as may intrinsic factors, such as lack of dNTPs, conflicts with the transcription machinery, and secondary DNA structures(7). It is now widely accepted that DNA replication stress is a main cause of genomic instability and a driver of tumorigenesis(8–11). For example, oncogenes can induce replication stress and genomic instability, by reducing dNTP levels available for replication or by increasing conflicts between the replication and transcription machinery(12,13). Consistent with this, tumor cells have higher basal replication stress than non-transformed cells(14–16).

Replication stress induces the replication stress checkpoint initiated by ATR and Chk1 kinases(17). These kinases coordinate the protection of replication forks, the inhibition of late-origin activation, the prevention of cell cycle progression, and both the induction of repair and the restarting of replication once the stress resolves. Fork protection can be achieved by fork slowdown, or by fork remodeling(18). One of the proposed models for fork remodeling under replication stress is fork reversal, a four-way junction structure (also called as “chicken foot”)(19,20), though other remodeling events also occur(21). Indeed, we have previously shown that, upon acute depletion of dNTP, some stalled forks do not regress but instead show nascent DNA disengagement from replisomes, which generates long stretches of parental single-stranded DNA (ssDNA)(22). Remodeled forks can restart DNA synthesis once the replication stress disappears, but if it persists, forks can collapse and DNA replication must be reinitiated by activating dormant origins or by recombination mechanisms, such as break-induced replication (BIR)(23,24).

Unsurprisingly, homologous recombination proteins engage in the replication stress response. Over recent years, diverse evidence has indicated that some homologous recombination proteins, such as BRCA2, Rad51, and Rad51 paralogs, participate in fork restart both after collapse and earlier in DNA protection and fork remodeling(21,25–28). During homologous recombination, Rad51 binds to the ssDNA generated by the nucleolytic processing of the DNA ends of a double-strand break (DSB). This cooperative binding generates a Rad51 nucleofilament that catalyzes homology recognition and DNA strand exchange(29). However, Rad51 has different roles under replication stress, independent from its strand exchange activity and even its ability to form nucleofilaments. First, it protects under-replicated parental ssDNA gaps to allow their post-replicative repair(30–32). Second, it induces fork reversal independent of the ability to form stable nucleofilaments, though a stable nucleofilament protects tracks of newly synthesized DNA in regressed forks from degradation by nucleases such as MRE11, EXO1, or DNA2(33–38). Finally, fork restart from stalled and collapsed forks appears to rely on strand exchange-dependent and -independent roles of Rad51(39–41). Consistent with the presence of basal replication stress in cancer cells and the role of Rad51 in dealing with replication stress, some cancer cells overexpress Rad51(42,43), making it a potential novel target for anti-cancer therapies(44–46).

We have previously shown that Rad51 is present at the replication forks of non-transformed human cells under both non-stress conditions and dNTP depletion induced by hydroxyurea (HU)(22), consistent with reports in yeast(31) and tumor cells(36,47). To better understand the relevance of this homologous recombination protein during the S phase, we analyzed the consequences of disrupting Rad51 during replication in non-transformed human cells and human colorectal cancer (CRC) cells, which have basal replication stress under control conditions. Our data show that neither Rad51 inhibition nor depletion affected fork stability or fork speed in non-transformed human cells under non-stress conditions, but that after recovery from an acute, or during a mild replication stress, efficient fork progression required Rad51 strand exchange activity. By contrast, Rad51 also sustained replication fork progression in CRC cells under control condition. This differential behavior reveals an opportunity for a novel therapeutic strategy that targets cancer cells specifically.

## Materials and Methods

### Cell lines and culture

All cell lines were purchased from the American Tissue and Cell Collection, ATCC. hTERT-RPE, hTERT-immortalized retinal pigment epithelial human cells (purchased from the American Tissue and Cell Collection, ATCC, Manassas, WV, USA), HCT116 (also from ATCC) and DLD-1 (purchased from Horizon Discovery Ldt., Cambridge, United Kingdom) both colorectal cancer human cells, were grown in Dulbecco’s modified Eagle’s medium (DMEM): HAM’s F12 (1:1) (Biological Industries, Beit HaEmek, Israel) supplemented with 6% fetal bovine serum (FBS, Biological Industries). All culture media were supplemented with 1% non-essential amino acids (Biological Industries), 2 mM L-glutamine (Sigma-Aldrich, St. Louis, MO, USA), 1 mM pyruvic acid (Sigma-Aldrich), 50 U/mL penicillin and 50 µg/mL streptomycin (both from Biological Industries).

### Drugs and cell synchronization

Drugs and their working concentrations were used as follows: 10mM HU in order to completely stall replication forks, 1mM HU in order to cause a mild replication stress or 0.1mM HU in order to cause a bearable replication stress; 25 µM B02 (Rad51 inhibitor) (Sigma-Aldrich); 25 µM 5-Chloro-2′-deoxyuridine (CldU) (Sigma-Aldrich); 250 µM 5-Iodo-2’-deoxyuridine (IdU) (Sigma-Aldrich); 50 µM 5-ethynyl-2′-deoxyuridine (EdU) (Invitrogen, Carlsbad, CA, USA); 10 µM 5-bromo-2’-deoxyuridine (BrdU) (Sigma-Aldrich); 250ng/mL nocodazole for hTERT-RPE (Sigma-Aldrich). Cell synchronization in S phase was performed by thymidine block as previously described(48).

### RNA interference

Transient siRNA experiments were performed using HiPerfect Transfection Reagent (Qiagen, Hilden, North Rhine-Westphalia, Germany). The following siRNA oligos were transfected at 50nM final concentration, using the manufacturer’s guidelines (Dharmacon, Lafayette, CO, USA): The siRNA sequences used were the following: a) Human ON-TARGETplus SMARTpools against Rad51 (Dharmacon, L-003530-00-005), which contains these 4 oligonucleotides: 5’-UAUCAUCGCCCAUGCAUCA-3’, 5’-CUAAUCAGGUGGUAGCUCA-3’, 5’-GCAGUGAUGUCCUGGAUAA-3’, 5’-CCAACGAUGUGAAGAAAUU-3’; b) Human ON-TARGETplus non-targeting pool (Dharmacon, D-001810-10-20) was used as a control containing these 4 oligonucleotides: 5’-UGGUUUACAUGUCGACUAA-3’, 5’-UGGUUUACAUGUUGUGUGA-3’, 5’-UGGUUUACAUGUUUU CUGA-3’, 5’-UGGUUUACAUGUUUUCCUA-3’.

### Western Blot (WB)

Cells were lysed with a buffer containing 67 mM Tris-HCl (pH 6.8) and 2% SDS. Denatured proteins were then resolved by SDS-PAGE and transferred to nitrocellulose membranes as previously described(49). Proteins were stained with Ponceau S (0.1% Ponceau reagent [Sigma, P3504] with 5% acetic acid). Incubation with primary antibodies was conducted overnight at 4°C. Antibodies against the indicated proteins were used as follows: Rad51 (Santa Cruz Biotechnology, Dallas, TX, USA, H-92, sc-8349; 1/200), Cyclin A2 (Santa Cruz Biotechnology, H-432 sc-751; 1/500), P-Chk1 S296 (Cell Signaling, Danvers, MA, USA, #2349; 1/1000), Lamin B (Santa Cruz Biotechnology, M-20 sc-6217; 1/200), GAP120 (Santa Cruz Biotechnology, sc-63; 1/100), CDK4 (Santa Cruz Biotechnology, sc-749; 1/500), Polymerase α (Santa Cruz Biotechnology, sc-5921; 1/200), Rad51 (Santa Cruz Biotechnology, sc-8349; 1/200). Proteins were visualized using EZ-ECL Kit (Biological Industries). Quantification was done by densitometry using Image Lab™ Software (Bio-Rad, Hercules, CA, USA, #1709690).

### Flow cytometry

Cells were harvested and fixed in 70% ethanol for at least 2 h at -20ºC before immunostaining and flow cytometry. Combined analysis of DNA content with propidium iodide (PI), BrdU (anti-BrdU, Abcam, Cambridge, United Kingdom, ab6326; 1/250) positive population and the MPM2 (anti-MPM2, Millipore, Burlington, MA, USA, #05-368; 1/250) positive population were performed with a BD FACSCalibur (Cytometry and Cell Sorting Core Facility, IDIBAPS) as previously described(49).

### 53BP1 immunofluorescence in hTERT-RPE

Previously grown attached in coverslips and treated, cells were rinsed with PBS and fixed in 2% PFA-containing PBS for 20 min at room temperature (RT). After several PBS washes, cells were permeabilized with 0.2% Triton X-100 in PBS for 10 min at RT and washed with PBS during 5 min. Cells were then incubated in blocking solution (3% FBS and 0.1% Triton X-100-containing PBS) for 1 h at RT. After blocking, cells were incubated with primary antibody (anti-53BP1; abcam, ab36823; 1/500) diluted in blocking solution during 45 min at 37°C. After 15 min washing in blocking solution at RT, cells were incubated with Alexa488-conjugated secondary antibody (Invitrogen; 1/500) diluted in blocking solution for 20 min at 37°C. Then, cells were washed again with blocking solution at RT and DNA was counterstained with DAPI (Sigma-Aldrich, D9564). For EdU staining, click reaction was performed previously to immunofluorescence during 30 minutes at RT with 1 µM Alexa488-azide (Invitrogen, A10266). Images were acquired in a Zeiss LSM880 confocal microscope and analysed using Fiji software.

### High Content Screening 53BP1 and γH2AX immunofluorescence

Cells were seeded, labelled with thymidine analogues (CldU and EdU) and treated in the µ-slide 8 well chamber (ibidi, Gräfelfing, Germany, 80826). After treatments and labelling, cells were permeabilized for 10 minutes at 4 ºC with 0.5% Triton X-100 and fixed with 4% Paraformaldehyde containing PBS (131 mM NaCl, 1.54 mM KH2PO4, 5.06 mM Na2HPO4) for 10 minutes at RT. To denature DNA, samples were treated with 2N HCl dissolved in PBS containing 0.1% (v/v) Triton X-100 for 30 min at RT. For EdU staining, click reaction was performed during 30 minutes at RT with 1 µM Alexa488-azide (Invitrogen, A10266). Then, cells were incubated for 1 hour at RT with primary antibodies against CldU (Abcam; ab6326) and 53BP1 (Abcam, ab36823; 1/500) or γH2AX (Merck Millipore, Burlington, MA, USA 05636;1/3000) diluted in filtered DMEM: HAM’s F12 (1:1) supplemented as indicated in “Cell lines and culture” and 5% BSA. After a wash with PBS-T (PBS + 0.01% Tween20) cells were incubated overnight at 4°C with secondary antibodies. Finally, mounting media with DAPI (ibidi,50011) was added. Images were obtained with Zeiss LSM880 confocal microscopies (Confocal Microscopy Unit Core Facility, University of Barcelona) with a PLAN APO 63 × oil immersion objective (numerical aperture 1.4) and analyzed using Fiji software. All data obtained were managed and analyzed with R studio software(50,51).

### DNA fiber assay

DNA fiber assay was performed in accordance with a protocol described in(41). Anti-BrdU antibodies were used for CldU (Abcam; ab6326; 1/1000) and IdU (Becton Dickinson, Franklin Lakes, NJ, USA; 347580; 1/200) labelling. Images were obtained using Leica TCS-SL or Zeiss LSM880 confocal microscopies (Confocal Microscopy Unit Core Facility, University of Barcelona) with a PLAN APO 63× oil immersion objective (numerical aperture 1.4), and then analysed using Fiji software. The number of fibers analysed in each experiment is indicated in the corresponding figure legend.

## RESULTS

### Rad51 activity maintains replication fork progression in non-transformed human cells upon an acute replication stress

Using the IPOND technique, we previously showed that Rad51 is present at active forks of non-transformed immortalized human (hTERT-RPE) cells under conditions of both unperturbed DNA synthesis and acute replication stress induced by HU treatment(48). Therefore, we first aimed to analyze whether Rad51 was needed to maintain replication fork progression under non-stress conditions. This involved treating non-transformed hTERT-RPE cells with B02, a small molecule that inhibits ssDNA-binding and strand exchange activities of Rad51(52), and measuring fork progression by DNA fiber analysis. As shown in Fig 1, treatment with B02 did not significantly affect replication fork progression (based on the length of the 5-iodo-2′-deoxyuridine (IdU)-labeled tracks), indicating that, despite the presence of Rad51 at replication forks, the maintenance of fork speed in hTERT-RPE cells under unperturbed conditions does not depend on Rad51 activity.

**Fig 1.**
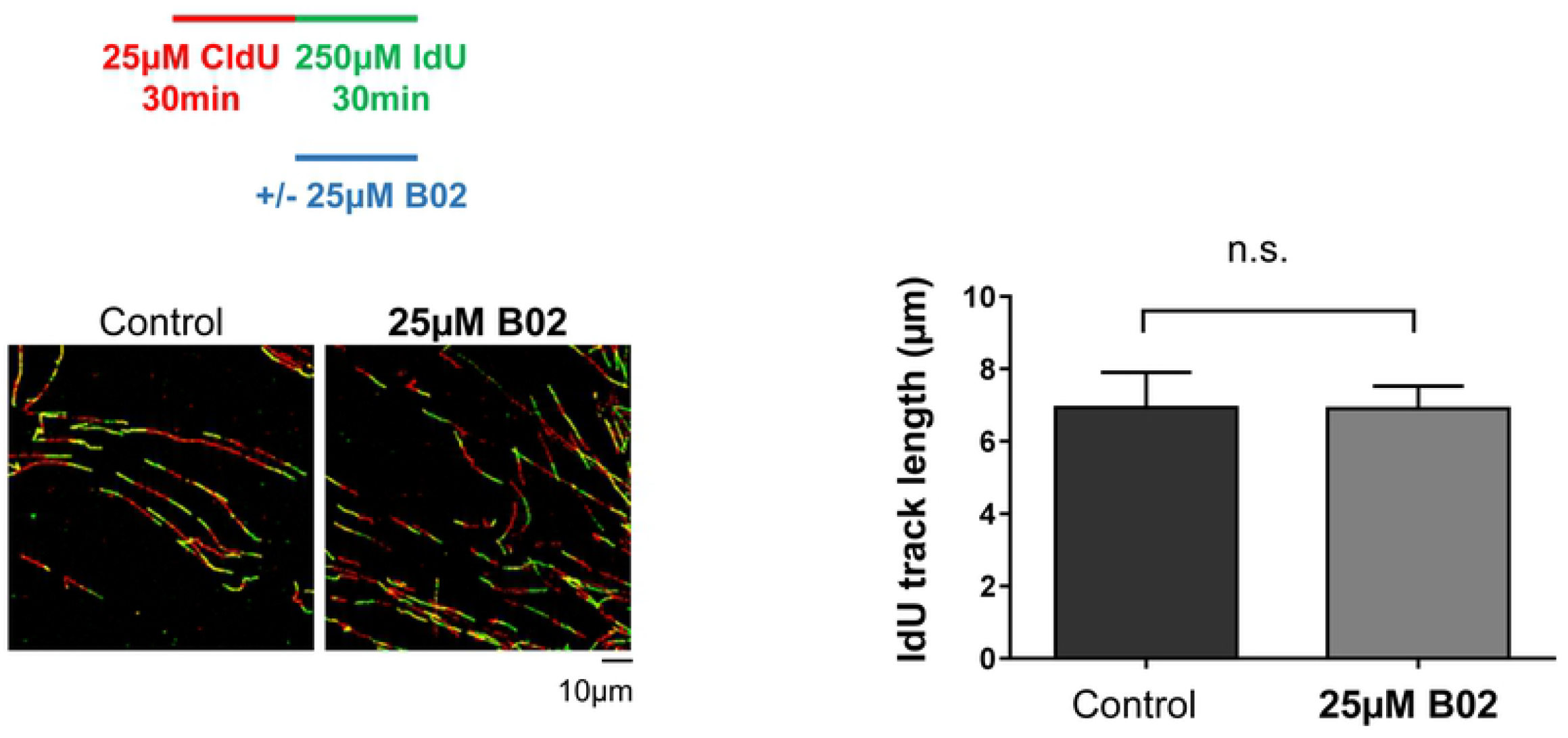
RAD51 inhibition does not affect fork progression under unperturbed conditions in hTERT-RPE. Cells were labelled as indicated (upper panel), adding the B02 inhibitor during the second analogue labelling. After that, cells were harvested and prepared for DNA fiber analysis. Representative images are shown (bottom-left panels). The IdU track length was measured. At least 200 fibers of each condition in each experiment were measured. Means and standard deviation (bars) of three experiments are shown (bottom-right panel, paired *t-*test, n.s.: non-statistically significant).

Next, we analyzed a possible role of Rad51 upon acute replication stress (10 mM HU treatment for 2 hours), which induced complete DNA replication arrest in hTERT-RPE cells without inducing detectable DNA damage (no increase in 53BP1/γH2Ax foci by immunofluorescence or in double-strand breaks by pulse-field gel electrophoresis)(22,48). As previously shown(48), we confirmed that most of the arrested replication forks could restart DNA synthesis after removing HU (Fig 2A). To ascertain that all the restarted forks constituted actual restarts and not the activations of dormant origins, which could be indistinguishable by the DNA fiber technique, we repeated the experiment with roscovitine (an inhibitor of cyclin-dependent kinases) that prevents new origin activation(53). As shown in Fig 2, even in the presence of roscovitine, upon HU removal almost all forks were able to reinitiate DNA synthesis and incorporate the second nucleotide analog. Cells were treated with B02 to analyze a putative role of Rad51 in fork restart. B02 was added at the same time as HU to inhibit Rad51 during fork arrest and was maintained during recovery. As shown in Fig 2A, we observed no significant differences in the percentage of forks able to restart DNA synthesis after HU removal, indicating that Rad51 is not essential to recover forks after an acute replication stress in hTERT-RPE cells. However, it remained possible that the presence of Rad51, rather than its activity, was necessary for fork restart, so we repeated the analysis by depleting cells of Rad51 using siRNA, but no differences in fork restart were observed (S1 Fig), indicating that restarting DNA synthesis from arrested forks did not strictly require the presence of Rad 51.

**Fig 2.**
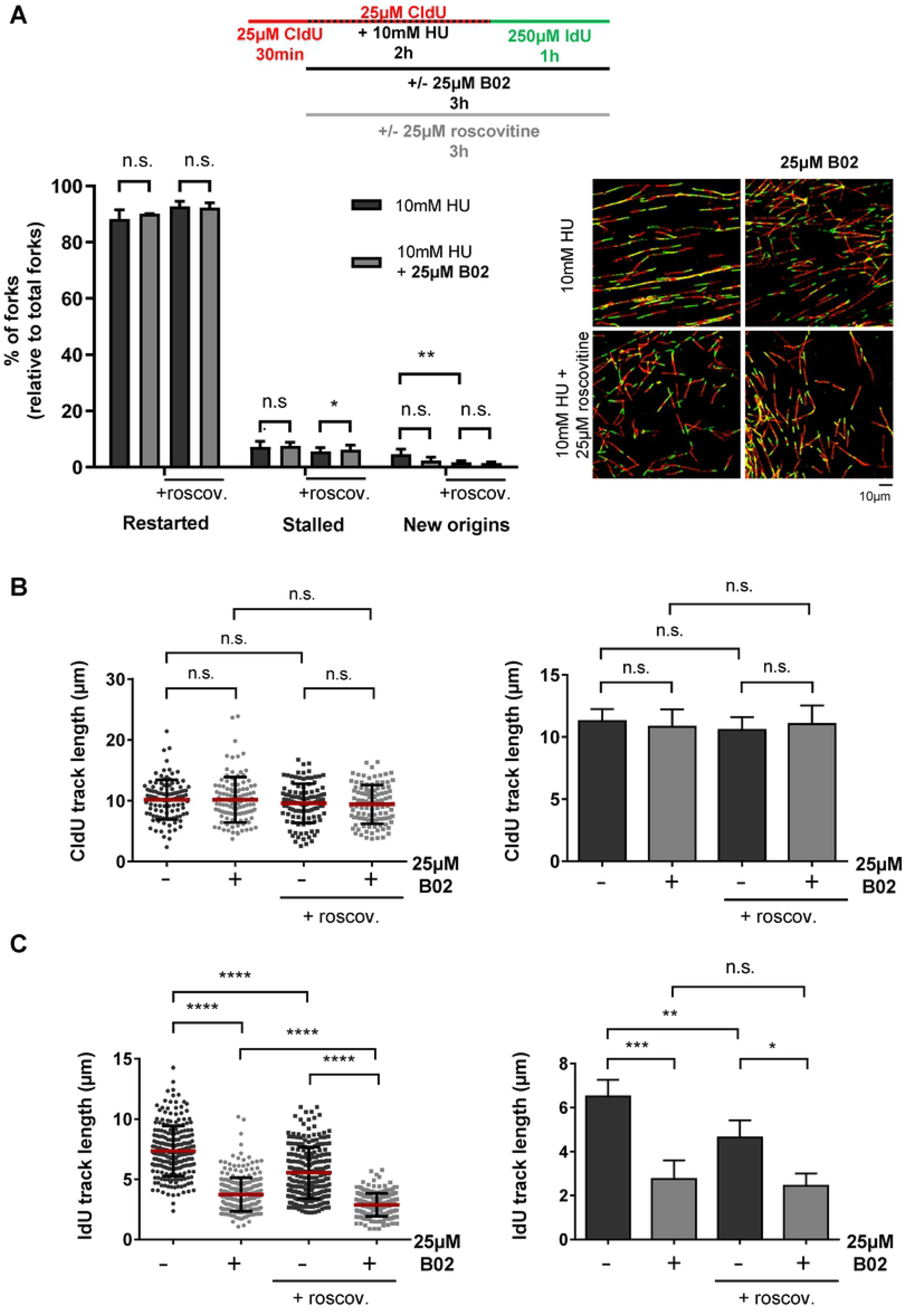
RAD51 inhibition does not affect the number of restarted forks neither causes fork degradation but impairs fork progression after an acute replication stress in hTERT-RPE cells. (**A**) Cells were labelled as indicated (upper panel), adding the B02 inhibitor and roscovitine with 10mM HU and the second analogue. After labelling, cells were harvested and prepared for DNA fiber analysis. At least 200 fibers of each condition in each experiment were used to calculate the percentage of restart, stalled forks and new origin firing events relative to total forks. Representative images are shown (bottom-right panel). Means and standard deviation (bars) of three experiments with (+roscov.) or without roscovitine are shown (bottom-left panel, paired t-test, n.s.: non-statistically significant, * P value < 0.05, ** P value < 0.01). (**B**) DNA fibers from (A) were used to measure CldU track length (first analogue). At least 300 fibers of each condition were measured. One representative experiment out of three is shown in left panel (Mann-Whitney test, n.s.: non-statistically significant; each dot represents a fiber track length). Means and standard deviation (bars) of three different experiments with (+roscov.) or without roscovitine are shown in right panel (paired t-test, n.s.: non-statistically significant). (**C**) DNA fibers from (A) were used to measure IdU track length (second analogue). At least 200 fibers of each condition in each experiment were measured. One representative experiment out of three is shown (bottom-left panel, Mann-Whitney test, **** P value < 0.0001). Means and standard deviation (bars) of three experiments with (+roscov.) or without roscovitine are shown (bottom-right panel, paired t-test, n.s.: non-statistically significant, * P value < 0.05, ** P value < 0.01, *** P value < 0.001).

Rad51 may protect nascent DNA from degradation after regressing arrested forks. We previously showed that no MRE11-dependent degradation of nascent DNA occurred during 2 h of HU treatment(22). Analysis of the length of DNA fibers labeled with 5-chloro-2′-deoxyuridine (CldU) (incorporated into DNA nucleotide before HU treatment) in cells treated with B02 (Fig 2B) or depleted of Rad51 (S2 Fig) show that Rad51 is not essential to protect nascent DNA in forks arrested by acute HU treatment. Interestingly, although the number of restarted forks was unaffected by B02 treatment, the images in Fig 2A highlight a defect in fork progression during restart in the setting of Rad51 inhibition. In fact, the measurement of DNA fiber tracks labeled with IdU (Fig 2C) revealed shorter lengths in B02-treated cells (independent of roscovitine), indicating that Rad51 activity facilitates restarted fork progression. This was also corroborated in cells depleted of Rad51 (S1D Fig).

By adding B02 during not only recovery but also fork arrest, these latter results could reflect the need for Rad51 activity during replication fork arrest to maintain forks in a conformation that allows efficient restart and progression or for the possibility that Rad51 activity directly participated in fork progression when they recovered from arrest. To test this, B02 was added 30 min after HU removal, which allowed Rad51 to perform its function during arrest and restart, but not during the progression of restarted replication forks. A recovery time of 30 minutes was chosen because this was the minimum time needed to detect nucleotide incorporation by fiber analysis after HU removal (data not shown). As shown in Fig 3, the length of newly synthesized DNA fiber tracks decreased under those conditions, indicating that Rad51 needed to be present continuously for progression after recovery from acute replication stress.

**Fig 3.**
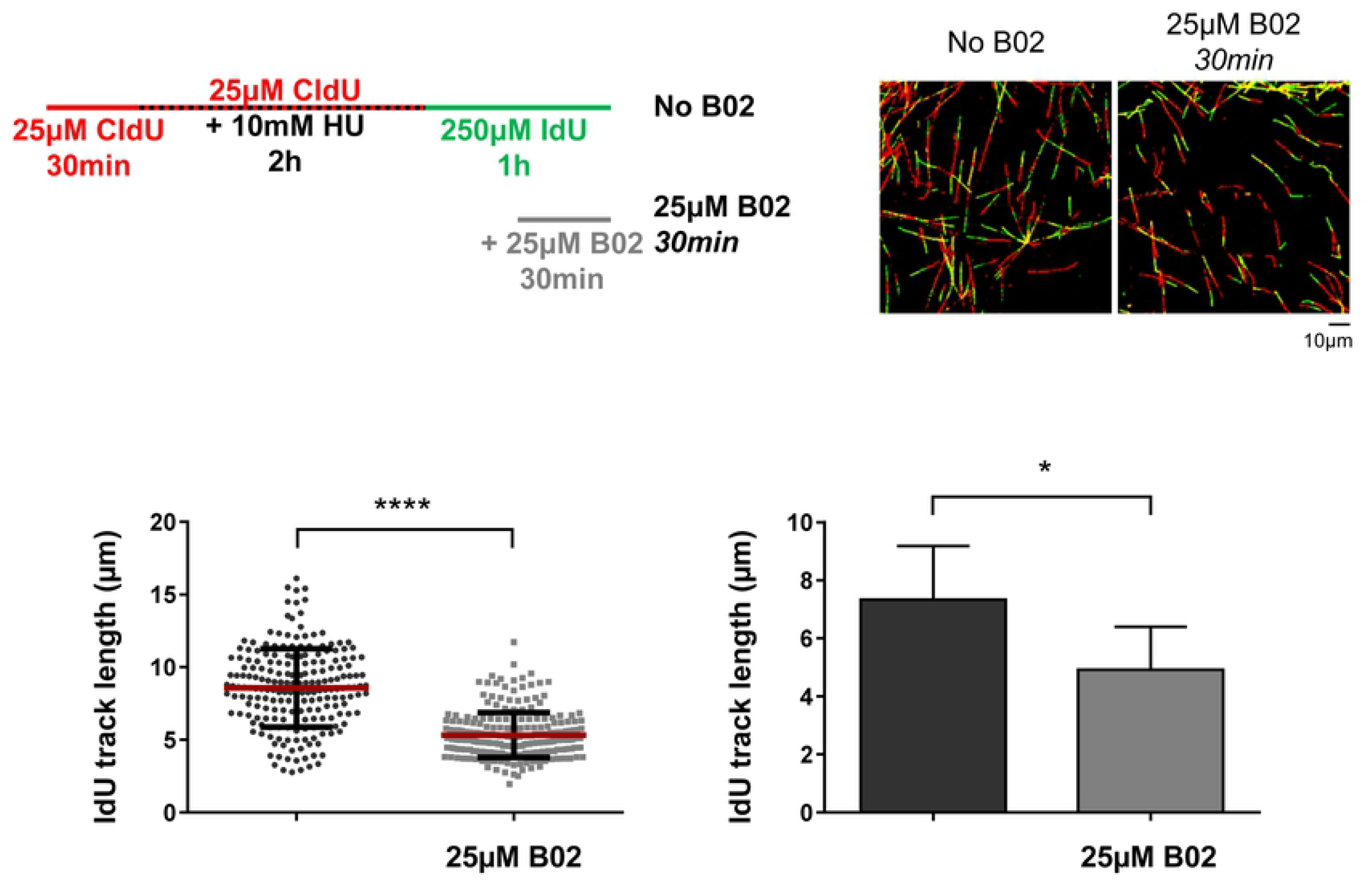
RAD51 is necessary for an efficient fork restart and progression after an acute replication stress in hTERT-RPE cells. Cells were labelled as indicated (upper-left panel), adding the B02 inhibitor after 30 minutes of HU release. After labelling, cells were harvested and prepared for DNA fiber analysis. Representative images are shown (upper-right panels). At least 200 fibers of each condition in each experiment were measured. One representative experiment out of three is shown in bottom-left panel (Mann-Whitney test, **** P value < 0.0001; each dot represents a fiber track length). Means and standard deviation (bars) of three experiments are shown in bottom-right panel (paired t-test, * P value < 0.05).

### Rad51 inhibition after acute replication stress induces DNA damage in non-transformed human cells

Although efficient progression of replication forks after reinitiation from acute HU-induced arrest required Rad51 activity, this did not prevent DNA replication from finishing. The same proportion of 5-bromo-2’-deoxyuridine (BrdU)-pulse labeled cells could progress into the G2/M phase 12 h after recovery from acute HU treatment, independent of Rad51 inhibition (S3 Fig). However, the proportion of cells arrested in G2 increased significantly in B02-treated cells when adding the B02 to cells during recovery from a 2h HU treatment-induced stress (Figs 4A and S3). To test if this G2 arrest correlated with increased DNA damage, we determined the percentage of cells with 53BP1 foci among all cells in S phase just before HU treatment (pulse-labeled with 5-ethyl-2’- deoxyuridine (EdU). Interestingly, our data showed a significant increase in DNA damage in cells that had been arrested in S phase for 2h by HU before B02 treatment (Figs 4B and S3). Rad51 depletion caused similar results, producing G2 arrest and an increase in cells with 53BP1 foci (S4 and S5 Figs). These results again support a key role of Rad51 following recovery from acute replication stress in hTERT-RPE cells.

**Fig 4.**
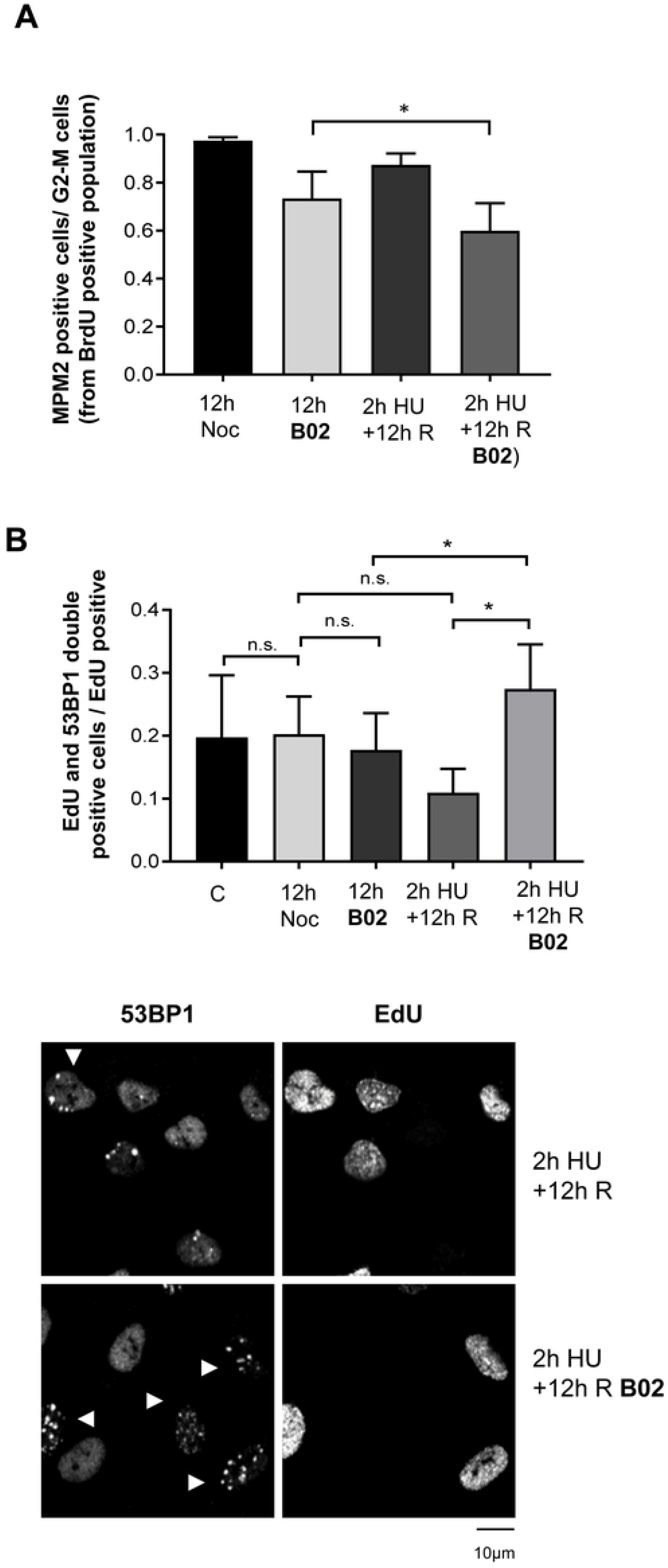
RAD51 inhibition affects mitotic entry, having more effect after an acute replication stress in hTERT-RPE cells. (**A**) Cells were labelled with BrdU and then treated during 2 hours with 10mM HU or left untreated .for 12 hours into nocodazole-containing fresh medium, without (12h Noc) or with RAD51 inhibitor (12h B02) (upper panel). HU treated cells were release for 12h in fresh media without (2h HU+12h R) or with B02 during the last 11h30’ (2hHU+12h R B02). Flow cytometry analysis of approximately 15000 cells was performed to analyse the S-phase population, initially labelled with BrdU analogue (BrdU-488 positive cells), after 12 hours. Cell cycle progression was analysed by measuring mitotic cells (MPM2-647 positive from BrdU-488-positive population) relative to cells into G2-M phases. Means and standard deviation (bars) of six experiments are shown (bottom panel, paired t-test, * P value < 0.05, ** P value < 0.01). (**B**) Cells were pulse-labelled with EdU analogue during 30 minutes (Control). Then, cells were treated as in (A) Finally, click reaction and 53BP1 immunofluorescence were performed. Representative images are shown (lower panels). At least 100 cells were counted for condition in each experiment. Means and standard deviation (bars) of three experiments in control and four experiments in other conditions are shown. The proportion of cells presenting both EdU and 53BP1 foci (more than six) relative to EdU positive cells is shown (upper panel, paired t-test, n.s.: non-statistically significant, * P value < 0.05, ** P value < 0.01).

### Rad51 inhibition reduces fork progression in non-transformed human cells under mild replication stress

Our data, using a 10mM HU treatment, support a role of Rad51 in cell recovery from replication stress causing complete fork arrest. Given that replication challenges encountered by cells due to intrinsic factors or chemotherapy might be milder and only slow the replication forks, we wanted to test whether Rad51 was essential for fork progression under these situations. To this end, cells were treated with either 1 mM HU (which induces a strong reduction in fork speed, but not a complete arrest, and an increase in P-Chk1(54); Fig 5A) or 0.1 mM HU (which also reduces fork speed but only causes a milder defect and does not increase P-Chk1; Fig 5A). Of note, B02 treatment of S phase cells did not increase P-Chk1 either when applied alone or in combination with HU (Fig 5A). The lengths of DNA tracks labeled with the second nucleotide analog indicated that Rad51 inhibition produced a significant reduction in nucleotide incorporation in the presence of both 1 mM HU (Fig 5B) and 0.1 mM HU (Fig 5C). These results strongly support the hypothesis that, even under very mild replication stress, Rad51 has an important role in maintaining fork progression.

**Fig 5.**
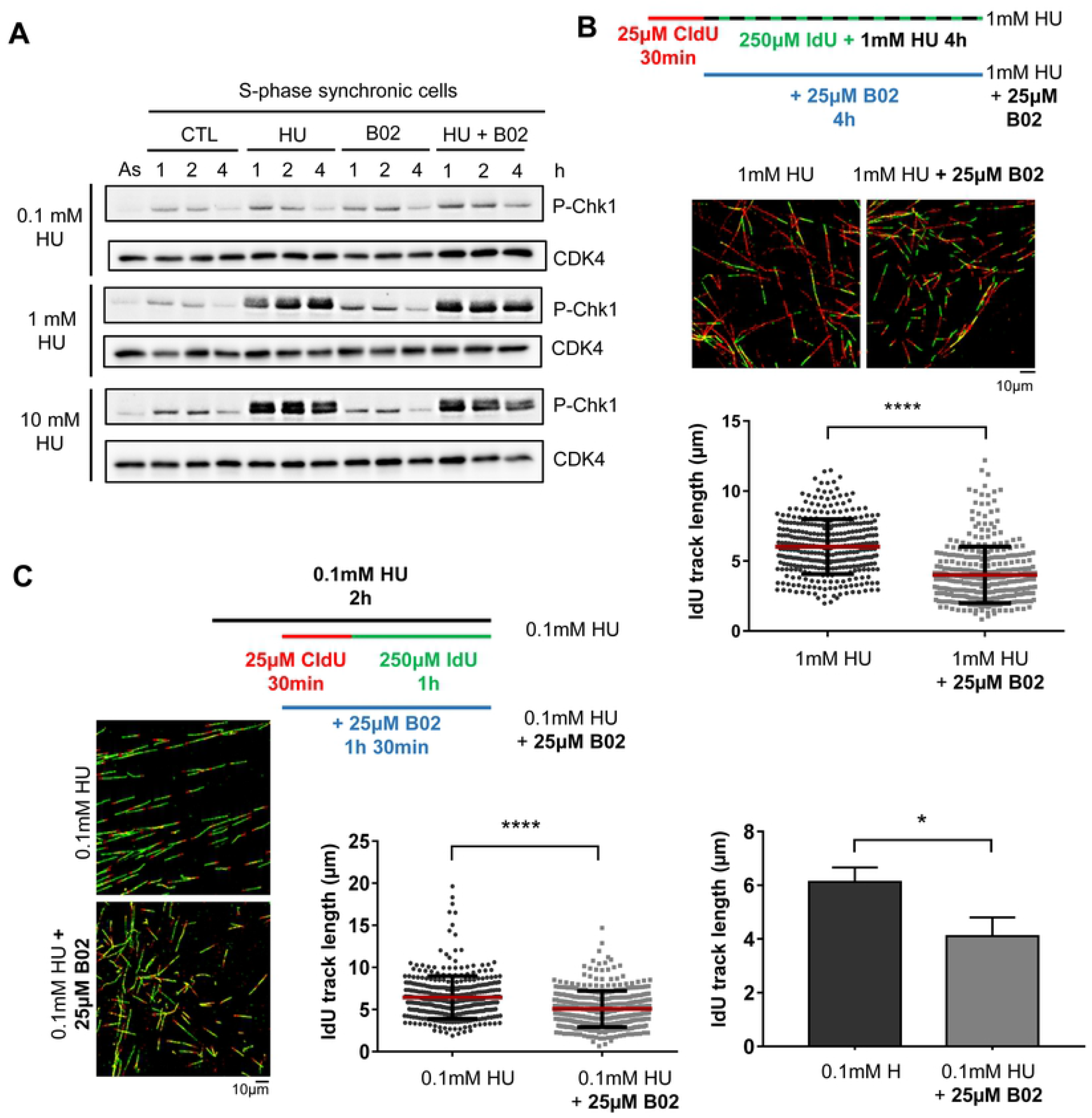
RAD51 inhibition affects fork progression under mild or bearable replication stress in hTERT-RPE cells. (**A**) Cells were synchronized in S phase by a thymidine block and 2h release. After that, they were treated with HU, B02 or left untreated (CTL) for the period indicated in the Fig. As.:asynchronic cells. Cells were lysed and P-Chk1 (Ser296) and CDK4 (loading control) were analyzed by western blot. (**B**) Cells were labelled as indicated (upper panel), adding the B02 inhibitor during the second analogue and HU treatment. After labelling, cells were harvested and prepared for DNA fiber analysis. Representative images are shown. At least 300 fibers of each condition in each experiment were measured. Quantification of a representative experiment out of two is shown (bottom-right panel, Mann-Whitney test, **** P value < 0.0001). (**C**) Cells were labelled as indicated (upper panel). HU was added 30 minutes before DNA labelling (this dose of HU is not completely inhibiting replication). B02 was added where it is indicated. After labelling, cells were harvested and prepared for DNA fiber analysis. Representative images are shown (left panels). At least 250 fibers of each condition in each experiment were measured. One representative experiment out of four is shown in middle graph (Mann-Whitney test, **** P value < 0.0001; each dot represents a fiber track length). Means and standard deviation (bars) of four experiments are shown right graph (paired t-test, * P value < 0.05).

### Rad51 inhibition reduces fork progression in cancer cells with basal replication stress

Transformed cells characteristically have higher basal replication stress that leads to an increase in P-Chk1 and a higher dependency of these cells on the DNA replication checkpoint(8). Nevertheless, replication forks in these cells manage to progress and complete DNA replication. This is shown by the HCT116 cell line derived from human colorectal adenocarcinoma. As previously shown(55), and corroborated in S6A Fig, these cells show higher levels of P-Chk1 under control conditions than non-transformed hTERT-RPE cells. Interestingly, these cancer cells also show higher levels of Rad51 than hTERT-RPE cells (S6A Fig). Having shown that cells under mild replication stress are dependent on Rad51 activity to maintain fork progression speed, we considered whether, in contrast to the findings in hTERT-RPE cells, Rad51 inhibition affected replication fork progression in HCT116 cells under control conditions. DNA fiber analysis showed that, while replication dynamics did not change under Rad51 inhibition (S6B Fig), the DNA track length of the second nucleotide decreased significantly in cells treated with B02 (Fig 6A). These results indicate that, contrasting with hTERT-RPE cells, HCT116 cells require Rad51 activity under control conditions to maintain replication fork speed. The HCT116 cells showed a prolonged S phase when treated with B02 (S6C Fig), which, interestingly, was corroborated with another CRC cell line (DLD-1) that also present basal replication stress(54) and higher levels of Rad51 (S6A Fig).

**Fig 6.**
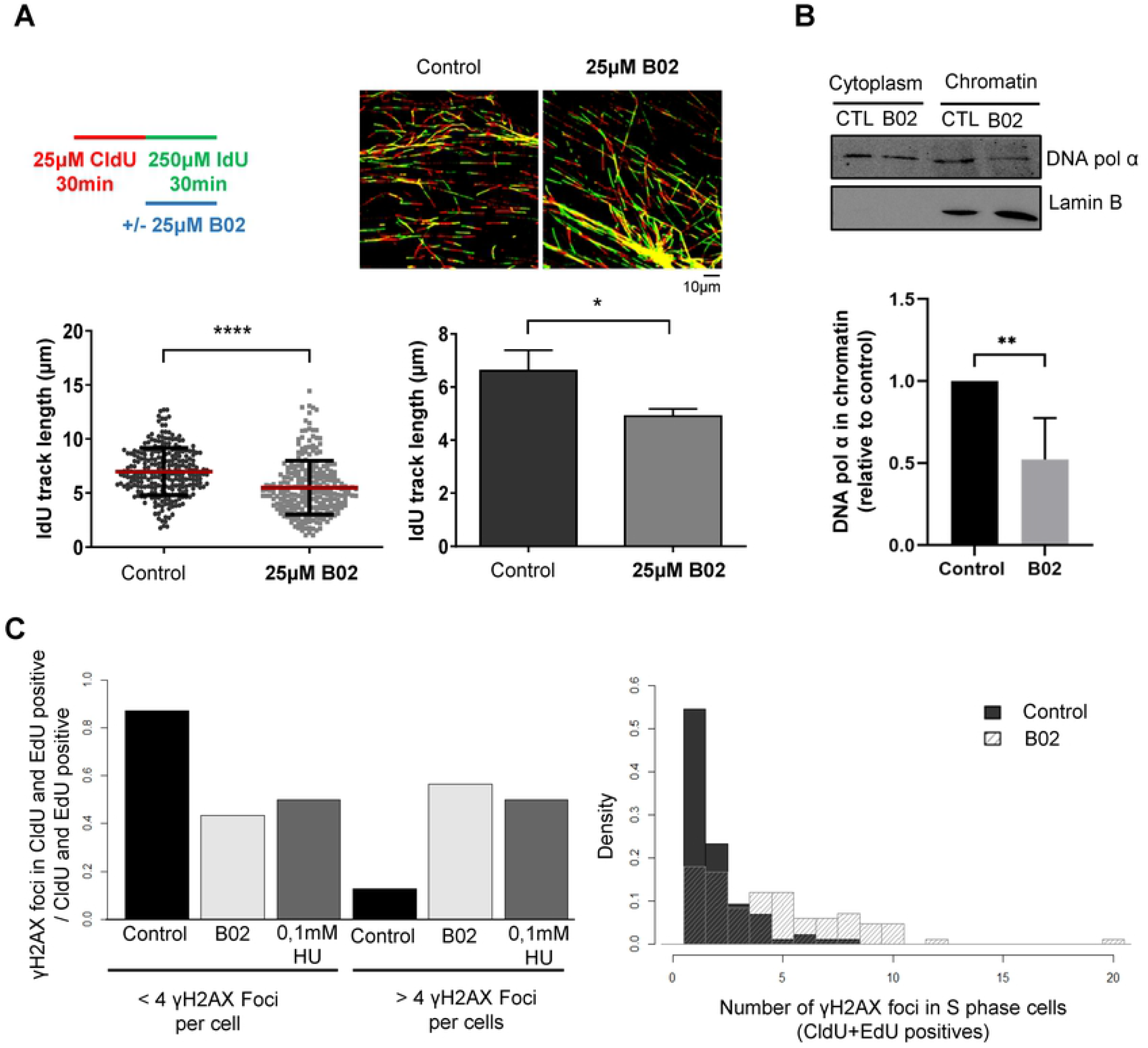
RAD51 inhibition affects fork progression during unperturbed conditions in HCT116 cells. (A) Cells were labelled as indicated (upper-left panel), adding the B02 inhibitor with the second analogue. After labelling, cells were harvested and prepared for DNA fiber analysis. Representative images are shown (upper-right panels). The IdU track length was measured. At least 250 fibers of each condition in each experiment were measured. One representative experiment out of three is shown in bottom-left panel (Mann-Whitney test, **** P value < 0.0001; each dot represents a fiber track length). Means and standard deviation (bars) of three experiments are shown (bottom-right panel, paired t-test, * P value < 0.05). (B) Cells were treated (B02) or not (CTL) with B02 for 4h and then cytoplasm and chromatin fraction purified as indicated in the methods section. DNA polymerase α (DNA pol α) and Lamin B (as control of chromatin purification and loading in the gel) were analyzed by western blot. A representative WB (upper panel) and quantification of 4 biological replicates (lower panel; mean and standard deviation, t-student test, ** P value< 0.01) are shown. (C) Cells were pulse labeled with CldU, treated with B02, HU, or no-drug (control) and pulse-labeled again with EdU. After processing for Immunofluorescence to detect CldU, EdU and γH2AX, images were acquired and analyzed as indicated in the methods section. A representative experiment is shown. B02 versus Control γH2AX foci number distribution are significantly different according to to Mann-Whitney test, **** P value < 0.0001.

Experiments performed in *Xenopus* egg extracts demonstrated that Rad51 interacts with DNA polymerase α to promote its association with stalled forks and prevent an increase in ssDNA gaps at replication forks (34). We therefore tested whether Rad51 could be involved in the recruitment of DNA polymerase α to the chromatin in HCT116 cells with a basal replication stress. Interestingly, HCT116 cells showed a slight reduction in chromatin-bound DNA polymerase α level under B02 treatment (Fig 6B). Of even greater interest, Rad51 inhibition in HCT116 cells during the S phase led to an increase in γH2Ax foci similar to that observed when these cells were treated with 0.1 mM HU (Figs 6C and S7A,). To analyze cells treated with B02 while replicating DNA, cells were pulse-labeled with CldU before and with EdU after a 4 h B02 treatment, and only cells labeled with both analogs were analyzed. By contrast, cells in S phase showed no increase in 53BP1 foci (S7B Fig), indicating that most probably ssDNA but not DSB is being generated.

## DISCUSSION

Replication fork progression during the S phase may encounter different impediments that cause fork slowing or stalling. These stalled forks require protection, remodeling, and reinitiation to avoid genome instability. Homologous recombination proteins, including BRCA1/2, Rad51, and Rad51 paralogs have emerged as key factors(19,25,28,56,57).

While most advances have been made using Xenopus egg extracts and human tumor cells (mainly U2OS cells), we analyzed the role of Rad51 in S phase progression in a non-transformed cell line, hTERT-RPE, comparing control conditions and different HU treatment conditions that do not induce fork collapse. Importantly, we mainly used B02 that inhibits the DNA strand exchange activity of human RAD51 and prevents Rad51 binding to ssDNA(52). Unlike using Rad51 knockdown with siRNA, this allowed us to study the effect of inhibiting Rad51 specifically during the S phase. In human non-transformed cells, Rad51 was found to be dispensable for fork progression under normal conditions while it helped to maintain replication fork progression under replications stress conditions. Consistent with the presence of basal replication stress in CRC tumor cells, these cells depended on Rad51 for proper DNA replication in unperturbed conditions.

In agreement with results using other normal cells, TD40 cells(58) or *Xenopus* extracts(30), we show that Rad51 is found in active replication forks interacting with nascent DNA under unperturbed DNA synthesis in non-transformed hTERT-RPE cells, but that it does not have an essential role in fork progression. Although we did not exclude the possibility that an absence of Rad51 produces very small ssDNA gaps (undetectable by DNA fiber analysis), as seen in *Xenopus* by electron microscopy, this should neither affect fork progression nor induce checkpoint activation during the S phase.

We previously showed that hTERT-RPE cells treated with 10 mM HU for 2 h displayed fork arrest, but that, contrasting with prolonged HU treatment(22,39,41,48), these stalled forks did not collapse because DNA synthesis restarted in almost 90% of them after removing HU (Fig 2). Since a dramatic increase in Rad51 has previously been observed in association with these arrested forks(22), we analyzed its role in nascent DNA protection, fork restart, and fork progression after restart. Other authors have shown that Rad51 nucleofilament formation on nascent DNA is essential to prevent DNA degradation in remodeled reversed forks; however, Rad51 (but not the nucleofilament formation) is also necessary for fork reversal and, consequently, degradation is observed in cells with non-functional BRCA1 but not in Rad51-depleted cells(26,35–37,56). Another fork remodeling mechanism that predominates after acute HU treatment in hTERT-RPE is the disengagement of nascent DNA from the replisome(22). We show here that Rad51 was not necessary to protect nascent DNA from degradation of these remodeled forks.

Concerning fork restart, different groups have used tumor and normal cells to demonstrate that Rad51-mediated strand invasion facilitates reversed fork restart, although both Rad51-dependent and Rad51-independent fork restart occurs in most cases(41). B02 did not affect the percentage of restarted forks after 2 h of 10 mM HU treatment, consistent with Rad51 having a mostly non-essential function in restarting stalled forks in non-transformed hTERT-RPE cells. The fact that fork reversion was not prevalent under our conditions could explain these differences; additionally, the presence of B02 during HU treatment could have further reduced the percentage of reversed forks.

It should be noted that, although forks managed to restart without Rad51 function in the first hour after HU removal, the resulting fork progression decreased significantly. This was observed not only when we added B02 during HU treatment but also when it was added at 30 min during recovery (when forks have already restarted). This indicates that progression after recovering from an acute replication stress requires the continued presence of Rad51, although it leaves the door open for the possibility that Rad51 may have a role in maintaining fork structure during arrest so that it can enable an efficient restart. Rad51 activity upon restarting stalled forks was important both to maintain fork speed and to avoid acquiring DNA damage, because when B02 was added during recovery from acute replication stress, the percentages of cells with more than six 53BP1 foci and of cells arrested in G2/M transition both increased. Costanzo’s group demonstrated, using *Xenopus* extracts, that Rad51 is essential for restarting collapsed forks (with DSB generated at the arrested forks by nucleases), and that absence of Rad51 produces gaps in the replicated DNA that are filled during G2(34). Our data add that Rad51 is essential to avoid generating DNA damage when synthesis restarts from non-collapsed arrested forks. Furthermore, we demonstrated the requirement for Rad51 not only in non-transformed cells to maintain progression speed upon restarting stalled forks but also under milder replication stress that reduces S phase progression without completely arresting the replication forks (1 mM or 0.1 mM HU treatment)(54).

Our results also prompted us to study if B02 treatment could inhibit fork progression in tumor cells with replication stress under control conditions. U2OS tumors cells have been extensively used to analyze the role of Rad51 during S phase with different inductors of replication stress: Rad51-mediated fork slowing has been described in response to BET inhibitors (which generate replication and transcription conflicts)(59), camptothecin, or mitomycin C(36). The authors propose that this is due to the role of Rad51 in inducing fork reversal, but no research has described a role of Rad51 in fork progression under unperturbed conditions in these tumor cells. We used CRC cells for three main reasons: (1) they present basal replication stress, as assessed by high levels of phosphorylated Chk1; (2) oncogenic KRAS present in HCT116 cells have increased Rad51 expression, which is a negative prognostic marker for colorectal adenocarcinoma(60), and (3) they are less sensitive to chemotherapy-induced DNA damage(61). Interestingly, B02 treatment of HCT116 cells reduced replication fork speed under basal conditions, which led to reduced S phase progression. These results were corroborated by other CRC cell lines (e.g., DLD-1) that also harbor an oncogenic KRAS allele.

Reduced replication fork progression induced by B02 in the presence of replication stress is consistent with the proposed role of Rad51 in re-priming DNA synthesis by facilitating DNA polymerase α to associate with chromatin(34). As demonstrated previously in *Xenopus* extracts, the present results showed that Rad51 also facilitates the interaction of DNA polymerase α and chromatin in HCT116 cells under unperturbed conditions, explaining the reduced fork speed observed with B02 treatment and the increase in γH2Ax foci observed in these cells.

Our data also explain why a BRCA2-derived peptide that targets Rad51 function elicited olaparib-induced cell death of U2OS human osteosarcoma cells but not of noncancerous cell lines(62). Tumor cells, with basal replication stress under unperturbed conditions, are more dependent on Rad51 activity than non-transformed cells for accurate DNA replication. This suggests that combining therapy with B02 and compounds that inhibit DNA repair, such as PARP inhibitors, could target increased tumor cell death.

## ACKNOWLEDGMENTS

This work was supported by grants from the Ministerio de Economia y Competitividad (SAF2016-76239-R; PID2019-105483RB-I00; for N.A.), an FI fellowship from the Generalitat de Catalunya (for S.F.), and an FPU fellowship from the Ministerio de Educación, Cultura y Deporte (for F.U.). We thank the personnel of the Advanced Microscopy Unit of CCiT-UB (Campus Clinic) for their help in setting up the image acquisition and analysis.

---

**S1 Fig. RAD51 depletion does not affect the number of restarted forks but impairs fork progression after an acute replication stress in hTERT-RPE cells**. (**A**) Cells were transfected with the indicated siRNA (NT: non-target) and 48 hours later cells were harvested for WB analysis with RAD51. Lamin B (LamB) was used as a loading control (upper panel). hTERT-RPE transfected cells were labelled as indicated (bottom panel). After labelling, cells were harvested and prepared for DNA fiber analysis. **(B)** Representative DNA fiber images are shown. **(C)** At least 200 fibers of each condition in each experiment were used to calculate the percentage of restart, stalled forks and new origin firing events relative to total forks. Means and standard deviation (bars) of three experiments are shown. The statistical analysis was performed just in HU-treated cells (paired t-test, non-statistically significant differences were found). (**D**) DNA fibers from were used to measure IdU track length (second analogue). At least 200 fibers of each condition in each experiment were measured. One representative experiment out of three is shown (bottom-left panel, Mann-Whitney test, **** P value < 0.0001).

**S2 Fig. RAD51 depletion does not cause fork degradation after an acute replication stress in hTERT-RPE cells**. hTERT-RPE cells were transfected with the indicated siRNA (NT: non-target) and 48 hours later cells were harvested for WB analysis with RAD51 antibody. Lamin B (LamB) was used as a loading control (upper-left panel). hTERT-RPE transfected cells were labelled as indicated (upper-right panel). After labelling, cells were harvested and prepared for DNA fiber analysis. Representative images are shown (bottom-right panels). The IdU track length was measured. At least 300 fibers of each condition in each experiment were measured. One representative experiment out of three is shown (bottom-left panel, Mann-Whitney test, n.s.: non-statistically significant).

**S3 Fig. RAD51 inhibition affects mitotic entry, although hTERT-RPE cells are able to arrive to G2 phase**. hTERT-RPE cells were labelled with BrdU and then treated during 2 hours with 10mM HU or left untreated for 12 hours into nocodazole-containing fresh medium, without (12h Noc) or with RAD51 inhibitor (12h B02) (upper panel). After HU treatment, cells were released into nocodazole-containing fresh medium, without (12h Noc) or with RAD51 inhibitor, added after 30 minutes of HU release (11h 30min B02). Flow cytometry analysis of approximately 15000 cells was performed to analyze the S-phase population, initially labelled with BrdU analogue (BrdU-488 positive cells). Cell cycle progression was analyzed by measuring mitotic cells (MPM2-647 positive from BrdU-488-positive population) relative to cells into G2-M phases (obtained by black DNA profiles from BrdU-488-positive population). Related to Figure 4.

**S4 Fig. RAD51 depletion does not impair replication recovery after an acute replication stress in hTERT-RPE cells**. hTERT-RPE cells were transfected with the indicated siRNA (NT: non-target) and 48h later cells were labelled with BrdU and then treated with 10mM HU or left untreated (12h release) into nocodazole-containing media for 12 hours. HU-treated cells were then released into nocodazole-containing fresh medium for 12 hours (2h HU + 12h release). Flow cytometry analysis of approximately 15000 cells was performed to analyse the S-phase arrested (BrdU-488 positive) cells after HU treatment, and the recovery from this stress measuring mitotic (MPM2-647 positive) cells from BrdU positive population.

**S5 Fig. RAD51 depletion increases genomic instability in hTERT-RPE cells**. hTERT-RPE cells were transfected with the indicated siRNA (NT: non-target) and 48 hours later cells were treated with 10mM HU for 2 hours or left untreated for 12 hours (Control). After HU treatment, cells were released into fresh medium for 12 hours. Finally, 53BP1 immunofluorescence was performed. The control for KD was shown (upper-left panel). Representative images from each condition are shown (upper-right panels). Two cells with more than six 53BP1 foci, indicated with a white arrowhead in the representative images from RAD51-depleted population, are shown in more detail (upper-right panels). At least 500 cells were counted for NT-depleted cells and 200 cells were counted for RAD51-depleted cells in each experiment. Means and standard deviation (bars) of percentage of cells presenting more than six 53BP1 foci of two experiments in control and three experiments in HU conditions are shown (bottom panel). The statistical analysis was performed just in HU-treated cells (unpaired t-test, ** P value < 0.01).

**S6 Fig. RAD51 inhibition does not affect replication dynamics but reduces S phase progression of CRC cells**. (**A**) Cells were lysed and P-Chk1 (Ser296) and Rad51 were analyzed by western blot. Ponceau was used as loading control (**B**) HTC116 cells were labelled as indicated in the upper-panel, adding the B02 inhibitor with the second analogue. After labelling, cells were harvested and prepared for DNA fiber analysis. Representative images are shown in Figure 6. DNA fibers were used to calculate the percentage of restart, stalled forks and new origin firing events relative to total forks. Around 1500 fibers from three independent experiments were counted in each condition. The average of those experiments is shown. Error bars represent standard deviation (paired *t*-test, n.s.: non-statistically significant). Related to Figure 6. (**C**) HCT116 cells were pulse labelled with BrdU and allow to proceed cell cycle in the absence of any drug (CTL, green profile) or in the presence of B02 (B02, red profile). Cells were harvested at the indicated times, fixed, permeabilized, and stained with propidium Iodide (PI) and with anti-BrdU antibody under denaturing conditions. DNA content (PI) of BrdU positive cells is shown. (**D**) as in (C) but with DLD1 cells.

**S7 Fig. Effect of B02 on γH2AX and 53BP1 foci in S phase HCT116 cells**. (**A**) Representative images of γH2AX, CldU and EdU immunofluorescence used for the quantification in Figure 6C are shown. (**B**) Cells were treated as in figure 6, but 53BP1 instead of γH2AX was analyzed by immunofluorescence. Related to figure 6.

## REFERENCES

1. Branzei D, Foiani M. Maintaining genome stability at the replication fork. Nat Rev Mol Cell Biol. 2010;11(3):208–19.

2. Kang S, Kang MS, Ryu E, Myung K. Eukaryotic DNA replication: Orchestrated action of multisubunit protein complexes. Mutat Res - Fundam Mol Mech Mutagen. 2018;809(March 2017):58–69.

3. Deegan TD, Diffley JFX. MCM: One ring to rule them all. Curr Opin Struct Biol. 2016;37:145–51.

4. Cortez D. Replication-Coupled DNA Repair. Mol Cell. 2019;74(5):866–76.

5. Liu W, Krishnamoorthy A, Zhao R, Cortez D. Two replication fork remodeling pathways generate nuclease substrates for distinct fork protection factors. Sci Adv. 2020;6:eabc3598.

6. Marians KJ. Lesion Bypass and the Reactivation of Stalled Replication Forks. Annu Rev Biochem. 2018;87(4):1–22.

7. Muñoz S, Méndez J. DNA replication stress: from molecular mechanisms to human disease. Chromosoma. 2016 Jan 21;

8. Macheret M, Halazonetis TD. DNA replication stress as a hallmark of cancer. Annu Rev Pathol. 2015 Jan;10:425–48.

9. Bester AC, Roniger M, Oren YS, Im MM, Sarni D, Chaoat M, et al. Nucleotide Deficiency Promotes Genomic Instability in Early Stages of Cancer Development. Cell. 2011 Apr 29;145(3):435–46.

10. Gaillard H, García-Muse T, Aguilera A. Replication stress and cancer. Nat Rev Cancer. 2015 Apr 24;15(5):276–89.

11. Burrell RA, McClelland SE, Endesfelder D, Groth P, Weller M-C, Shaikh N, et al. Replication stress links structural and numerical cancer chromosomal instability. Nature. 2013 Feb 28;494(7438):492–6.

12. Primo LMF, Teixeira LK. Dna replication stress: Oncogenes in the spotlight. Genet Mol Biol. 2020;43(1):1–14.

13. Aird KM, Zhang G, Li H, Tu Z, Bitler BG, Garipov A, et al. Suppression of Nucleotide Metabolism Underlies the Establishment and Maintenance of Oncogene-Induced Senescence. Cell Rep. 2013;3(4):1252–65.

14. Boyer A-S, Walter D, Sørensen CS. DNA replication and cancer: From dysfunctional replication origin activities to therapeutic opportunities. Semin Cancer Biol. 2016 Jan 12;

15. Ubhi T, Brown GW. Exploiting DNA Replication Stress for Cancer Treatment. Cancer Res. 2019;78:1–11.

16. Nieto-Soler M, Morgado-Palacin I, Lafarga V, Lecona E, Murga M, Callen E, et al. Efficacy of ATR inhibitors as single agents in Ewing sarcoma. Oncotarget. 2016;7(37):58759–67.

17. Branzei D, Foiani M. The checkpoint response to replication stress. DNA Repair (Amst). 2009 Sep 2;8(9):1038–46.

18. Zeman MK, Cimprich KA. Causes and consequences of replication stress. Nat Cell Biol. 2014 Jan;16(1):2–9.

19. Quinet A, Lemaçon D, Vindigni A. Replication Fork Reversal: Players and Guardians. Mol Cell. 2017;68(5):830–3.

20. Neelsen KJ, Lopes M. Replication fork reversal in eukaryotes: from dead end to dynamic response. Nat Rev Mol Cell Biol. 2015 Feb 25;16(4):207–20.

21. Berti M, Teloni F, Mijic S, Ursich S, Fuchs J, Palumbieri MD, et al. Sequential role of RAD51 paralog complexes in replication fork remodeling and restart. Nat Commun. 2020;11(1).

22. Ercilla A, Feu S, Aranda S, Llopis A, Brynjólfsdóttir SH, Sørensen CS, et al. Acute hydroxyurea-induced replication blockade results in replisome components disengagement from nascent DNA without causing fork collapse. Cell Mol Life Sci. 2020;77(4):735–49.

23. Sakofsky CJ, Ayyar S, Malkova A. Break-induced replication and genome stability. Vol. 2, Biomolecules. 2012. p. 483–504.

24. Costantino L, Sotiriou SK, Rantala JK, Magin S, Mladenov E, Helleday T, et al. Break-induced replication repair of damaged forks induces genomic duplications in human cells. Science. 2014 Jan 3;343(6166):88–91.

25. Somyajit K, Saxena S, Babu S, Mishra A, Nagaraju G. Mammalian RAD51 paralogs protect nascent DNA at stalled forks and mediate replication restart. Nucleic Acids Res. 2015;gkv880.

26. Bhat KP, Cortez D. RPA and RAD51: Fork reversal, fork protection, and genome stability. Nat Struct Mol Biol. 2018;25(6):446–53.

27. Kolinjivadi AM, Sannino V, de Antoni A, Técher H, Baldi G, Costanzo V. Moonlighting at replication forks: a new life for homologous recombination proteins BRCA1, BRCA2 and RAD51. FEBS Lett. 2017 Jan 12;

28. Saxena S, Dixit S, Somyajit K, Nagaraju G. ATR Signaling Uncouples the Role of RAD51 Paralogs in Homologous Recombination and Replication Stress Response. Cell Rep. 2019;29(3):551–559.e4.

29. San Filippo J, Sung P, Klein H. Mechanism of eukaryotic homologous recombination. Annu Rev Biochem. 2008;77:229–57.

30. Hashimoto Y, Ray Chaudhuri A, Lopes M, Costanzo V. Rad51 protects nascent DNA from Mre11-dependent degradation and promotes continuous DNA synthesis. Nat Struct Mol Biol. 2010 Nov;17(11):1305–11.

31. González-Prieto R, Muñoz-Cabello AM, Cabello-Lobato MJ, Prado F. Rad51 replication fork recruitment is required for DNA damage tolerance. EMBO J. 2013;32(9):1307–21.

32. Aiello FA, Palma A, Malacaria E, Zheng L, Campbell JL, Shen B, et al. RAD51 and mitotic function of mus81 are essential for recovery from low-dose of camptothecin in the absence of the WRN exonuclease. Nucleic Acids Res. 2019;47(13):6796–810.

33. Zadorozhny K, Sannino V, Beláň O, Mlčoušková J, Špírek M, Costanzo V, et al. Fanconi-Anemia-Associated Mutations Destabilize RAD51 Filaments and Impair Replication Fork Protection. Cell Rep. 2017;21(2):333–40.

34. Kolinjivadi AM, Sannino V, De Antoni A, Zadorozhny K, Kilkenny M, Técher H, et al. Smarcal1-Mediated Fork Reversal Triggers Mre11-Dependent Degradation of Nascent DNA in the Absence of Brca2 and Stable Rad51 Nucleofilaments. Mol Cell. 2017;67(5):867–881.e7.

35. Bhat KP, Krishnamoorthy A, Dungrawala H, Garcin EB, Modesti M, Cortez D. RADX Modulates RAD51 Activity to Control Replication Fork Protection. Cell Rep. 2018;24(3):538–45.

36. Zellweger R, Dalcher D, Mutreja K, Berti M, Schmid JA, Herrador R, et al. Rad51-mediated replication fork reversal is a global response to genotoxic treatments in human cells. J Cell Biol. 2015;208(5):563–79.

37. Mijic S, Zellweger R, Chappidi N, Berti M, Jacobs K, Mutreja K, et al. Replication fork reversal triggers fork degradation in BRCA2-defective cells. Nat Commun. 2017;8(1):1–11.

38. Schlacher K, Wu H, Jasin M. A Distinct Replication Fork Protection Pathway Connects Fanconi Anemia Tumor Suppressors to RAD51-BRCA1/2. Cancer Cell. 2012 Jul 10;22(1):106–16.

39. Mason J, Chan Y-L, Weichselbaum RW, Bishop DK. Non-enzymatic roles of human RAD51 at stalled replication forks. Nat Commun. 2019;

40. Hashimoto Y, Puddu F, Costanzo V. RAD51-and MRE11-dependent reassembly of uncoupled CMG helicase complex at collapsed replication forks. Nat Struct Mol Biol. 2012;19(1):17–24.

41. Petermann E, Orta ML, Issaeva N, Schultz N, Helleday T. Hydroxyurea-stalled replication forks become progressively inactivated and require two different RAD51-mediated pathways for restart and repair. Mol Cell. 2010 Feb 26;37(4):492–502.

42. Raderschall E, Stout K, Freier S, Suckow V, Schweiger S, Haaf T. Elevated levels of Rad51 recombination protein in tumor cells. Cancer Res. 2002;62(1):219–25.

43. Maacke H, Opitz S, Jost K, Hamdorf W, Henning W, Krüger S, et al. Over-expression of wild-type Rad51 correlates with histological grading of invasive ductal breast cancer. Int J Cancer. 2000;88(6):907–13.

44. Alagpulinsa DA, Ayyadevara S, Shmookler Reis RJ. A small molecule inhibitor of RAD51 reduces homologous recombination and sensitizes multiple myeloma cells to doxorubicin. Front Oncol. 2014;4(OCT):1–11.

45. Huang F, Mazin A V. A small molecule inhibitor of human RAD51 potentiates breast cancer cell killing by therapeutic agents in mouse xenografts. PLoS One. 2014;9(6):e100993.

46. King HO, Brend T, Payne HL, Wright A, Ward TA, Patel K, et al. RAD51 Is a Selective DNA Repair Target to Radiosensitize Glioma Stem Cells. Stem Cell Reports. 2017;8(1):125–39.

47. Chaudhuri AR, Callen E, Ding X, Gogola E, Duarte AA, Lee JE, et al. Replication fork stability confers chemoresistance in BRCA-deficient cells. Nature. 2016;535(7612):382–7.

48. Ercilla A, Llopis A, Feu S, Aranda S, Ernfors P, Freire R, et al. New origin firing is inhibited by APC/CCdh1activation in S-phase after severe replication stress. Nucleic Acids Res. 2016;44(10):4745–62.

49. Llopis A, Salvador N, Ercilla A, Guaita-Esteruelas S, Barrantes I del B, Gupta J, et al. The stress-activated protein kinases p38α/β and JNK1/2 cooperate with Chk1 to inhibit mitotic entry upon DNA replication arrest. Cell Cycle. 2012 Oct 1;11(19):3627–37.

50. Rstudio Team. RStudio: Integrated development for R. RStudio, Inc., Boston MA. RStudio. 2019.

51. Team RC. R: A Language and Environment for Statistical Computing. Vienna, Austria. 2019;

52. Huang F, Mazina OM, Zentner IJ, Cocklin S, Mazin A V. Inhibition of homologous recombination in human cells by targeting RAD51 recombinase. J Med Chem. 2012;55(7):3011–20.

53. Meijer L, Borgne A, Mulner O, Chong JPJ, Blow JJ, Inagaki N, et al. Biochemical and cellular effects of roscovitine, a potent and selective inhibitor of the cyclin-dependent kinases cdc2, cdk2 and cdk5. Eur J Biochem. 1997 Jan;243(1–2):527–36.

54. Feu S, Unzueta F, Llopis A, Semple JI, Ercilla A, Guaita-Esteruelas S, et al. OZF is a Claspin-interacting protein essential to maintain the replication fork progression rate under replication stress. FASEB J. 2020;34(5):6907–19.

55. Bianco JN, Bergoglio V, Lin Y-L, Pillaire M-J, Schmitz A-L, Gilhodes J, et al. Overexpression of Claspin and Timeless protects cancer cells from replication stress in a checkpoint-independent manner. Nat Commun. 2019;10(1):910.

56. Schlacher K, Christ N, Siaud N, Egashira A, Wu H, Jasin M. Double-strand break repair-independent role for BRCA2 in blocking stalled replication fork degradation by MRE11. Cell. 2011 May 13;145(4):529–42.

57. Guerra C, Schuhmacher AJ, Cañamero M, Grippo PJ, Verdaguer L, Pérez-Gallego L, et al. Chronic pancreatitis is essential for induction of pancreatic ductal adenocarcinoma by K-Ras oncogenes in adult mice. Cancer Cell. 2007 Mar;11(3):291–302.

58. Su X, Bernal JA, Venkitaraman AR. Cell-cycle coordination between DNA replication and recombination revealed by a vertebrate N-end rule degron-Rad51. Nat Struct Mol Biol. 2008;15(10):1049–58.

59. Bowry A, Piberger AL, Rojas P, Saponaro M, Petermann E. BET Inhibition Induces HEXIM1-and RAD51-Dependent Conflicts between Transcription and Replication. Cell Rep. 2018;25(8):2061–2069.e4.

60. Tennstedt P, Fresow R, Simon R, Marx A, Terracciano L, Petersen C, et al. RAD51 overexpression is a negative prognostic marker for colorectal adenocarcinoma. Int J Cancer. 2013;132(9):2118–26.

61. Kalimutho M, Bain AL, Mukherjee B, Nag P, Nanayakkara DM, Harten SK, et al. Enhanced dependency of KRAS mutant colorectal cancer cells on RAD51-dependent homologous recombination repair identified from genetic interactions in Saccharomyces cerevisiae. Mol Oncol. 2017 Feb 7;

62. Trenner A, Godau J, Sartori AA. A Short BRCA2-derived cell-penetrating peptide targets RAD51 function and confers hypersensitivity toward PARP inhibition. Mol Cancer Ther. 2018;17(7):1392–404.

